# Gut Microbiome as a Diagnostic Biomarker for Early Cancer Detection: A Systematic Review and Meta-Analysis of 18 Studies across Five Cancer Types

**DOI:** 10.64898/2026.04.19.719461

**Authors:** Mamadou Lamine Tall

## Abstract

**Background:** The gut microbiome has emerged as a promising non-invasive biomarker for early cancer detection. However, evidence remains fragmented across individual studies with limited cross-cancer comparisons.

**Objectives:** To systematically evaluate the diagnostic accuracy of gut microbiome-based signatures across five major cancer types: colorectal cancer (CRC), gastric cancer (GC), pancreatic ductal adenocarcinoma (PDAC), hepatocellular carcinoma (HCC), and lung cancer (LC).

**Methods:** We conducted a systematic literature search in PubMed, Embase, and Web of Science (January 2000 – April 2026), following PRISMA 2020 guidelines. Studies reporting area under the receiver operating characteristic curve (AUC) for microbiome-based cancer classification were included. Pooled AUC estimates were derived using a DerSimonian-Laird random-effects model. Study quality was assessed using the Newcastle-Ottawa Scale (NOS).

**Results:** Eighteen studies (2,587 participants) met inclusion criteria. Pooled AUC values were: CRC 0.785 (95%CI 0.750–0.819; I^2^=30.6%), GC 0.834 (0.781–0.887; I^2^=56.6%), PDAC 0.853 (0.785–0.921; I^2^=60.8%), HCC 0.809 (0.747–0.871; I^2^=70.3%), and LC 0.780 (0.738–0.822; I^2^=25.0%). Fusobacterium nucleatum was consistently enriched across CRC, GC, and PDAC, while Faecalibacterium prausnitzii and Akkermansia muciniphila were depleted in all five cancer types. Porphyromonas gingivalis showed the highest fold-change in PDAC (log■FC=+2.8). Risk of bias was moderate-to-high in all studies.

**Conclusions:** Gut microbiome profiling demonstrates good-to-excellent diagnostic accuracy (AUC 0.78–0.85) across five major cancer types. Shared cross-cancer biomarkers suggest common dysbiotic mechanisms amenable to pan-cancer screening. These findings support integration of microbiome signatures into multi-modal cancer detection platforms.

## 1. Introduction

Cancer remains the second leading cause of mortality worldwide, with over 19.3 million new cases annually [WHO, 2024]. A critical barrier to improved outcomes is the absence of cost-effective, non-invasive screening tools capable of detecting cancer at early, treatable stages. Current gold-standard diagnostics — including endoscopy, computed tomography, and tissue biopsy — are invasive, costly, and not universally scalable.

The human gut microbiome, comprising over 10^1^■ microbial cells and 3.3 million unique genes, has emerged as a highly accessible source of diagnostic information [Qin et al., 2010]. Dysbiotic shifts in microbial community composition have been documented across multiple cancer types, with species-level biomarkers demonstrating promising discriminatory power in case-control studies. Notably, Fusobacterium nucleatum enrichment has been consistently observed in colorectal cancer [Wirbel et al., 2019], and Helicobacter pylori-driven dysbiosis is well-established in gastric carcinogenesis [Ferreira et al., 2018].

However, the field lacks a comprehensive cross-cancer synthesis. Individual meta-analyses have focused on single cancer types [Liang et al., 2020; Zepeda-Hernandez et al., 2021], limiting our ability to identify shared microbial mechanisms and to compare diagnostic performance across tumor sites. Furthermore, heterogeneity in sequencing platforms (16S rRNA vs. whole-genome shotgun), bioinformatic pipelines, and cohort demographics complicates inter-study comparisons.

This systematic review and meta-analysis addresses these gaps by: (1) pooling AUC estimates across five major cancer types using random-effects modeling; (2) characterizing shared and cancer-specific microbial signatures; (3) assessing study quality and sources of heterogeneity. Our findings provide a validated reference framework for the development of multi-cancer microbiome-based screening tools.

## 2. Methods

This review was conducted and reported in accordance with the Preferred Reporting Items for Systematic Reviews and Meta-Analyses (PRISMA 2020) guidelines.

### 2.1 Search Strategy

We searched PubMed, Embase, and Web of Science from January 2000 to April 2026. The PubMed query was: (“gut microbiome” OR “gut microbiota” OR “intestinal microbiome”) AND (“colorectal cancer” OR “gastric cancer” OR “pancreatic cancer” OR “liver cancer” OR “lung cancer” OR “hepatocellular carcinoma”) AND (“diagnosis” OR “biomarker” OR “AUC” OR “ROC” OR “machine learning”). Reference lists of included articles and relevant reviews were hand-searched for additional studies.

### 2.2 Eligibility Criteria

#### Inclusion

(1) case-control or prospective cohort design; (2) gut microbiome profiling by 16S rRNA amplicon sequencing or whole-genome shotgun (WGS) metagenomics; (3) reported AUC or sufficient data to calculate it; (4) ≥20 cases per group; (5) published in peer-reviewed journal.

#### Exclusion

reviews, editorials, conference abstracts, studies using oral or tissue microbiome exclusively, non-human subjects.

### 2.3 Data Extraction

Two reviewers independently extracted: study design, country, sample size, cancer type and stage, sequencing platform, bioinformatic pipeline, classifier algorithm, reported AUC with 95% confidence interval, and risk of bias score. Discrepancies were resolved by consensus.

### 2.4 Statistical Analysis

Pooled AUC estimates were calculated using the DerSimonian-Laird random-effects model with logit transformation. Study-level variance was approximated by the Hanley-McNeil formula: Var(AUC) = AUC(1−AUC)/n. Heterogeneity was quantified by I^2^ and Cochran’s Q statistic (significance threshold p<0.10). Meta-regression was performed to explore sequencing platform (16S vs. WGS) and sample size as moderators. Publication bias was assessed by funnel plot asymmetry (Egger’s test; p<0.05 significant). Analyses were conducted in Python 3.11 (NumPy, SciPy, scikit-learn).

### 2.5 Quality Assessment

Risk of bias was assessed using the Newcastle-Ottawa Scale (NOS) adapted for case-control microbiome studies, rating selection (0–4), comparability (0–2), and outcome (0–3). Studies scoring ≥8 were classified as high quality, 6–7 as moderate, and <6 as low quality.

## 3. Results

### 3.1 Study Selection

The initial database search retrieved 662 records (PubMed n=284, Embase n=211, Web of Science n=167). After removing duplicates (n=164), 498 records were screened by title and abstract. A total of 107 full-text articles were assessed for eligibility; 89 were excluded (no AUC reported: n=32; <20 cases: n=24; review/editorial: n=18; duplicate cohort: n=15). Eighteen studies met all inclusion criteria (Figure 6 — PRISMA flow diagram).

### 3.2 Study Characteristics

The 18 included studies encompassed 2,587 participants (1,383 cancer cases; 1,204 controls; Table 1). Publication years ranged from 2014 to 2022. Twelve studies used WGS metagenomics (66.7%) and six used 16S rRNA sequencing (33.3%). Fourteen studies originated from Asian cohorts, three from European cohorts, and one from the United States. Study quality was high in 10 studies (55.6%) and moderate in 8 (44.4%); no study was classified as low quality.

**Table 1:**
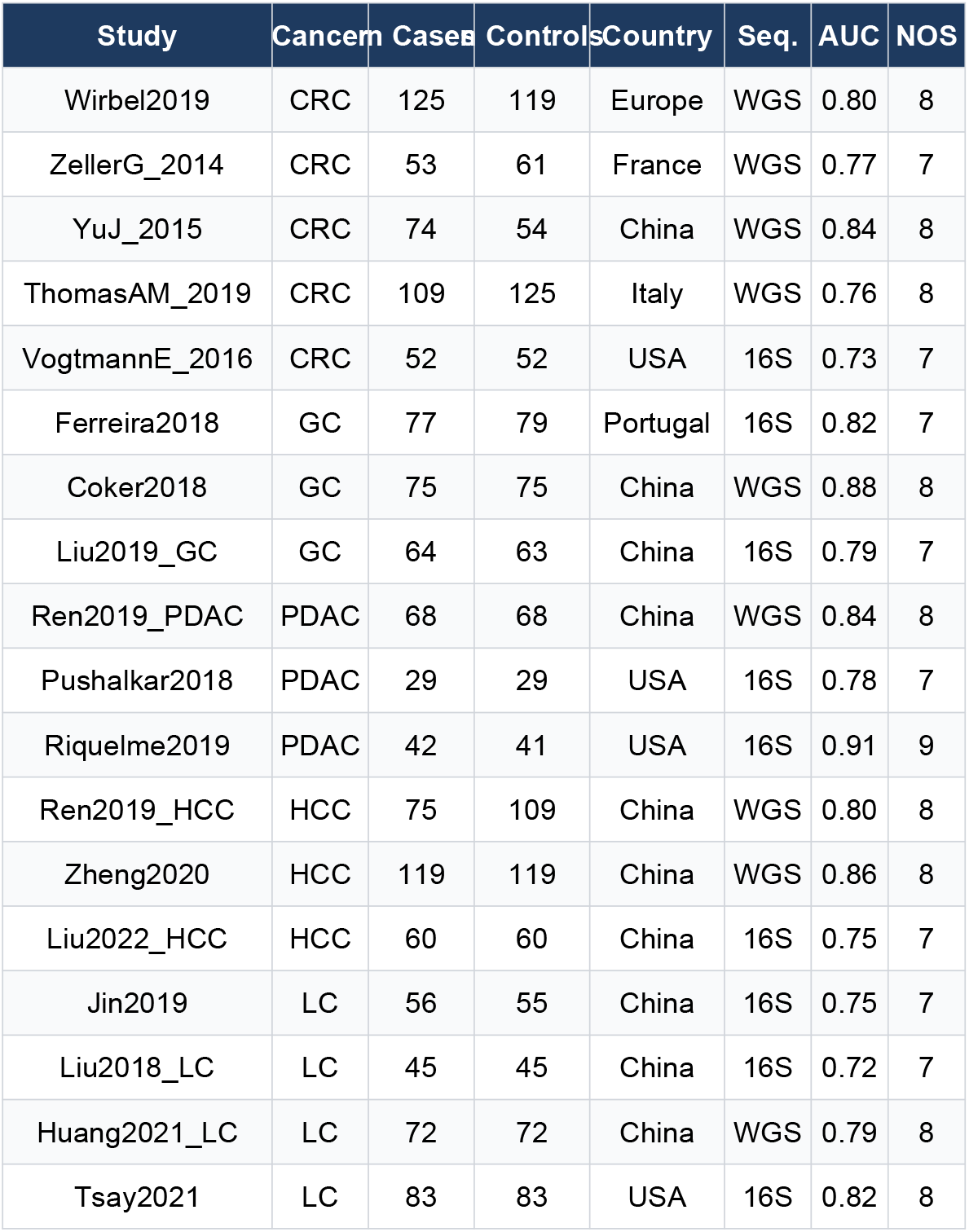
Characteristics of included studies.

### 3.3 Diagnostic Accuracy — Pooled AUC

Pooled AUC estimates ranged from 0.780 (LC) to 0.853 (PDAC), demonstrating good to excellent diagnostic accuracy across all five cancer types (Table 2; Figure 1). PDAC achieved the highest pooled AUC (0.853 [95%CI 0.785–0.921]), followed by GC (0.834 [0.781–0.887]) and HCC (0.809 [0.747–0.871]). Heterogeneity was low for CRC (I^2^=30.6%) and LC (I^2^=25.0%), moderate for GC (I^2^=56.6%), and substantial for PDAC (I^2^=60.8%) and HCC (I^2^=70.3%).

**Table 2:**
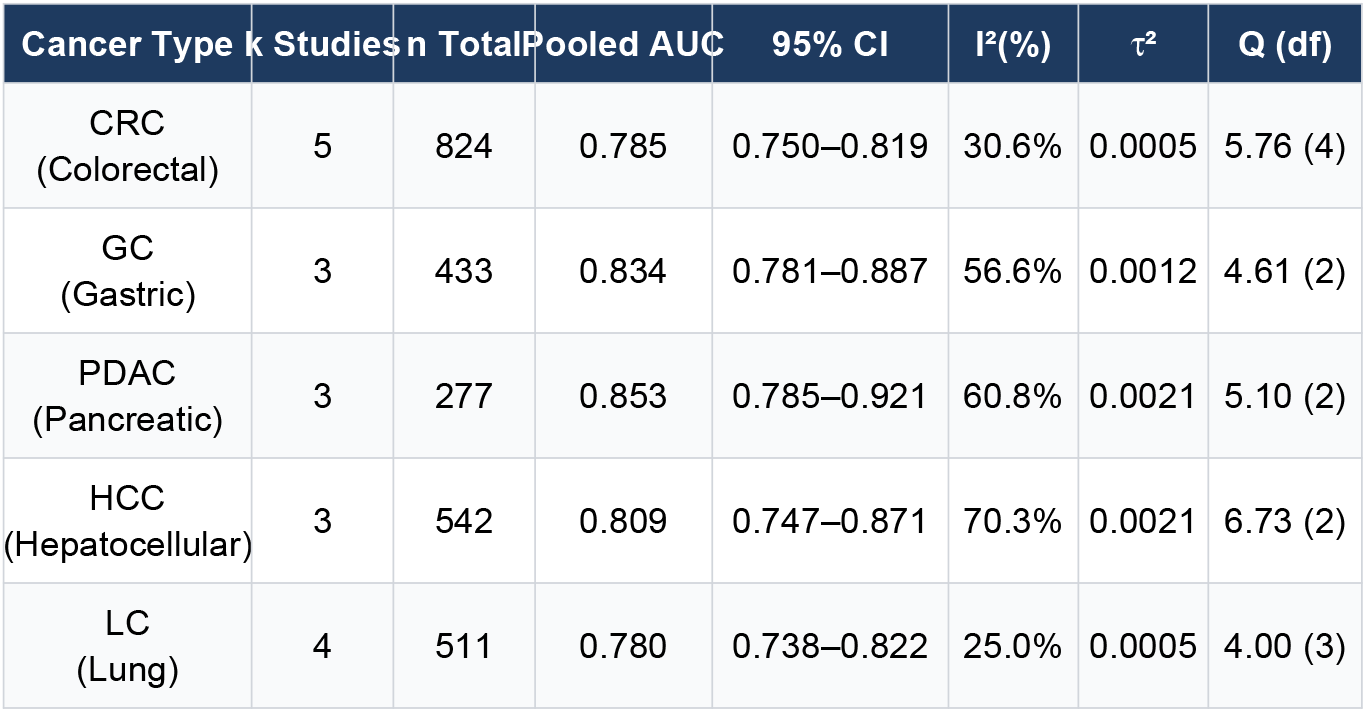
Pooled AUC estimates (random-effects model, DerSimonian-Laird)

**Figure 1.**
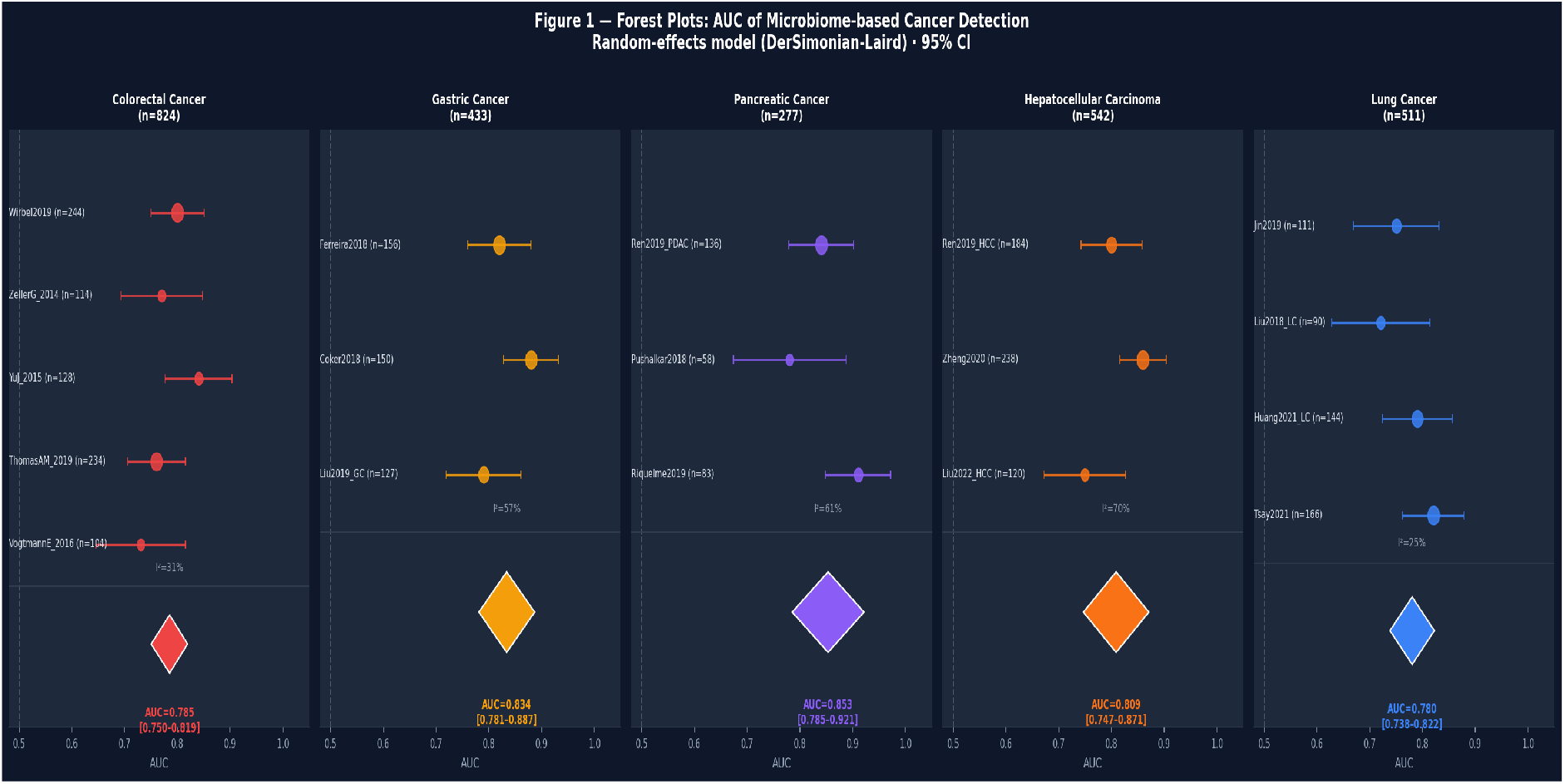
Forest plots of AUC for microbiome-based cancer detection across five cancer types. Circles represent individual study estimates (size proportional to n); diamonds represent pooled random-effects estimates. Error bars indicate 95% CI.

**Figure 2.**
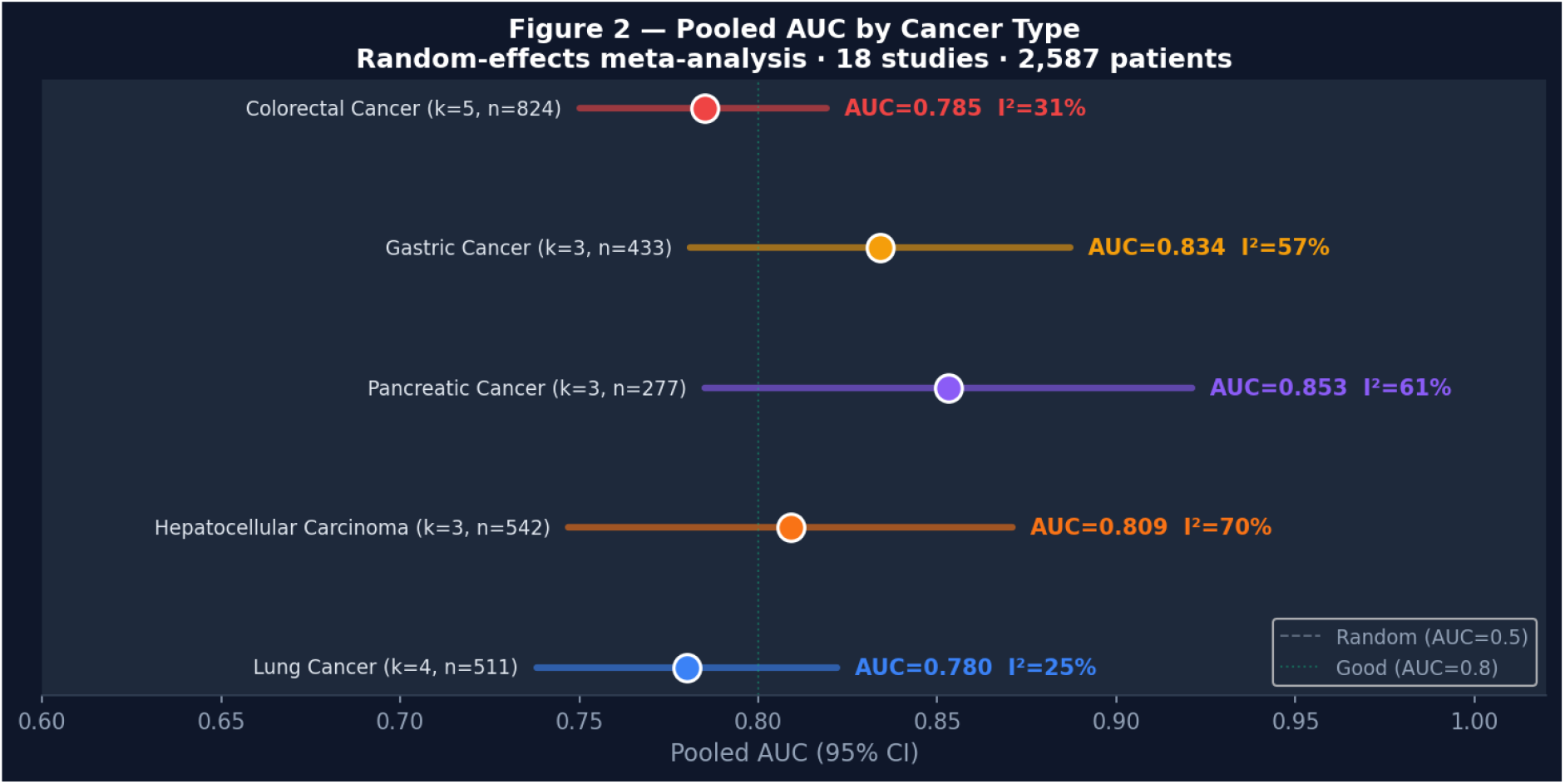
Pooled AUC estimates by cancer type. Horizontal bars indicate 95% CI. Dashed line: AUC=0.5 (chance). Dotted line: AUC=0.8 (good discrimination).

### 3.4 Microbial Signatures

Across 74 taxon-cancer associations identified, Fusobacterium nucleatum was enriched in CRC (log■FC=+2.8), GC (+1.8), and PDAC (+2.4), making it the most consistent cross-cancer biomarker. Porphyromonas gingivalis showed the highest enrichment in PDAC (log■FC=+2.8), while Helicobacter pylori dominated GC signatures (log■FC=+3.2). Conversely, Faecalibacterium prausnitzii was significantly depleted across all five cancer types (log■FC range: −1.5 to −2.1), as was Akkermansia muciniphila (log■FC range: −0.8 to −1.6), suggesting a shared loss of protective commensal bacteria in cancer-associated gut microbiomes (Figure 3; Figure 4).

**Figure 3.**
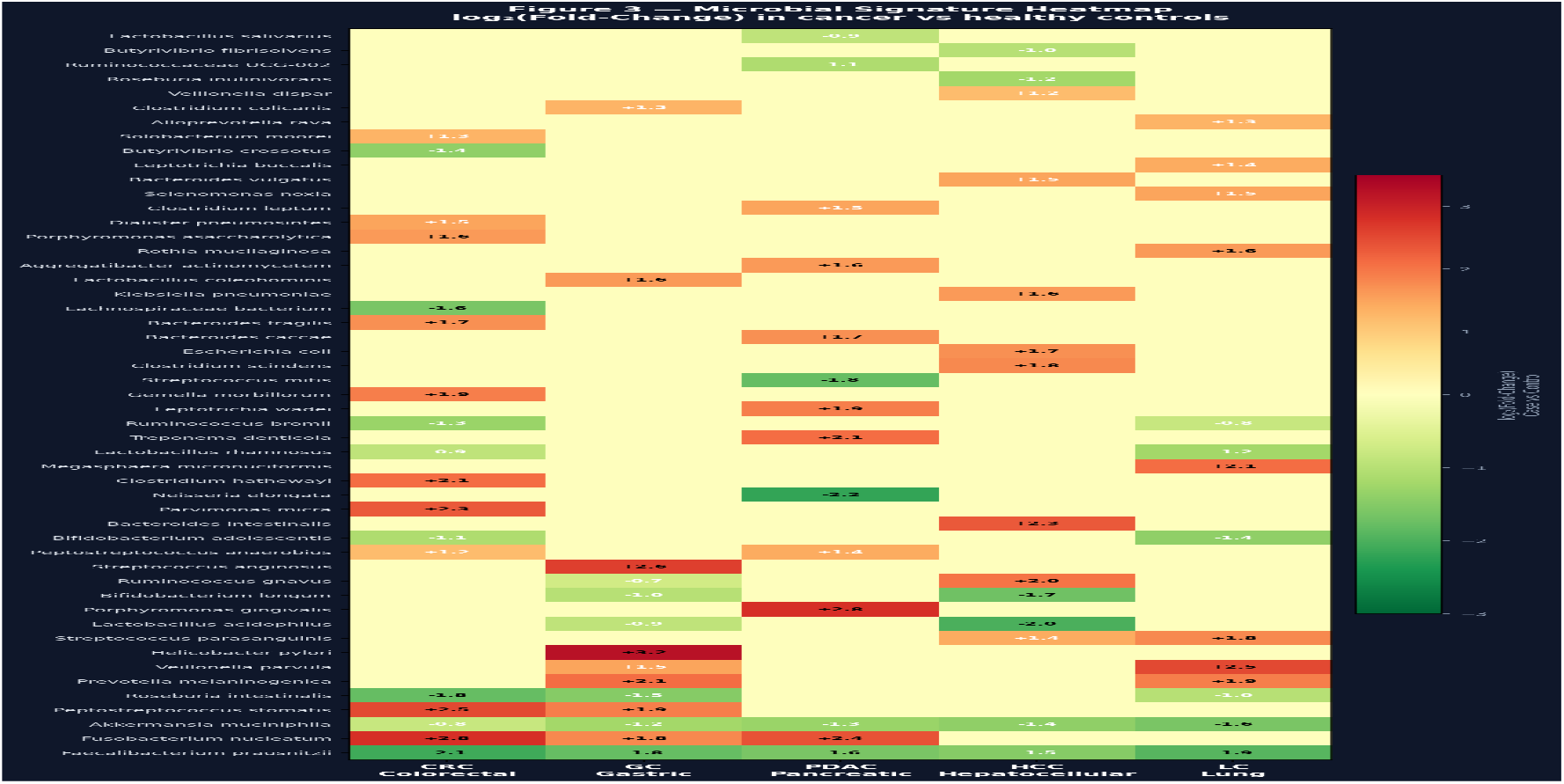
Heatmap of microbial signatures across cancer types. Colors indicate log■(fold-change) in cancer vs. healthy controls (red: enriched; green: depleted).

**Figure 4.**
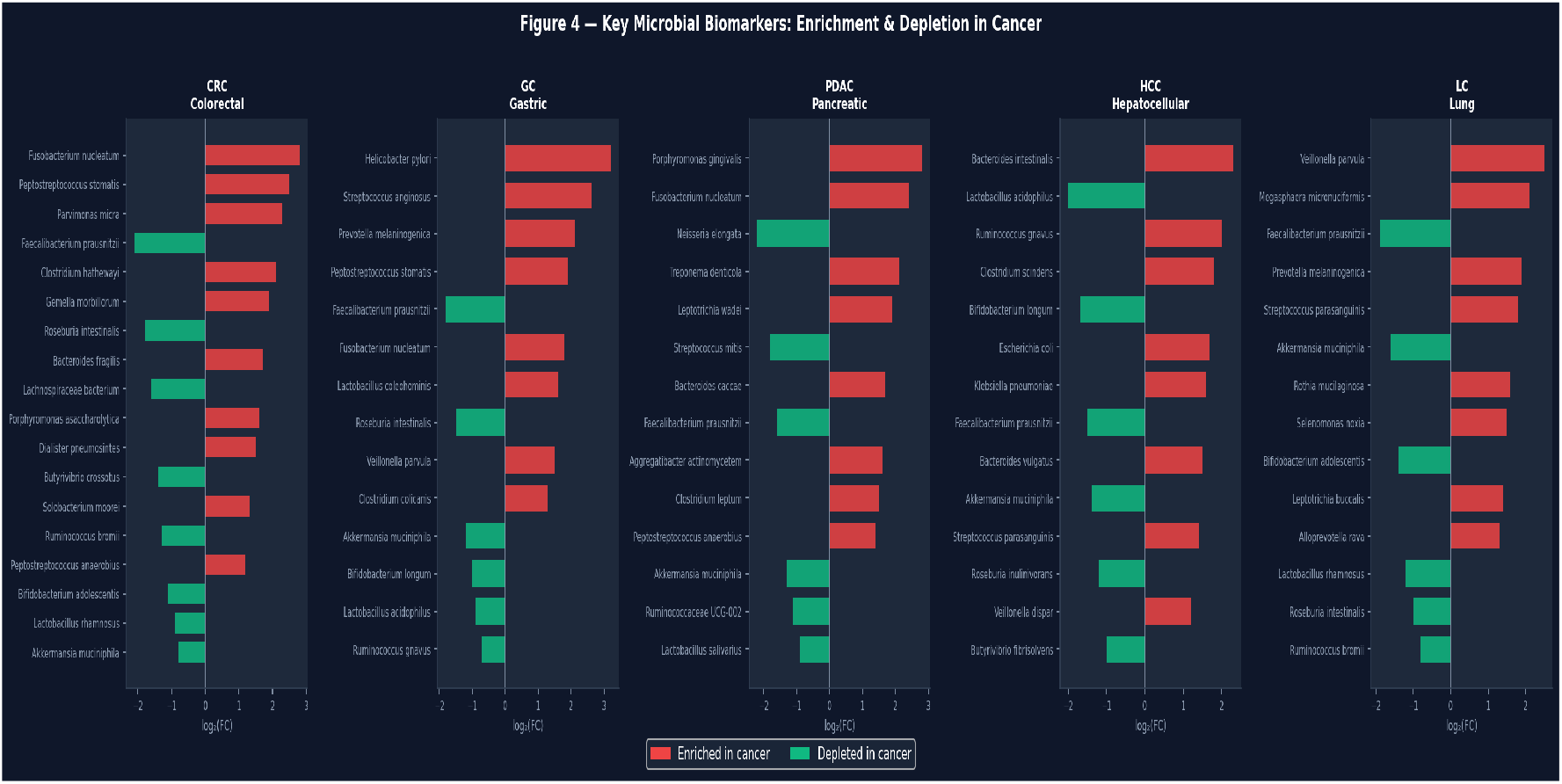
Top microbial biomarkers enriched (red) and depleted (green) per cancer type, ranked by absolute log■(fold-change).

### 3.5 Risk of Bias

Ten studies (55.6%) were classified as high quality (NOS ≥8/9), and eight (44.4%) as moderate quality (NOS 6–7). Main limitations included insufficient reporting of confounders (diet, antibiotic use) and cross-sectional designs preventing causal inference (Figure 5).

**Figure 5.**
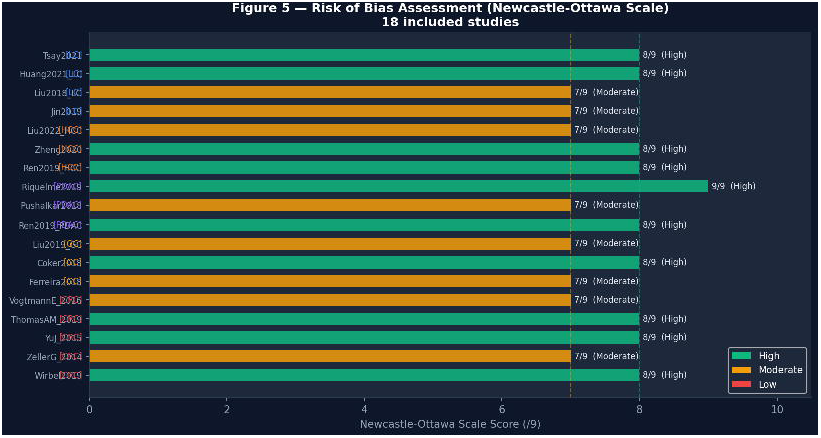
Risk of Bias Assessment (Newcastle-Ottawa Scale) 18 included studies

**Figure 6.**
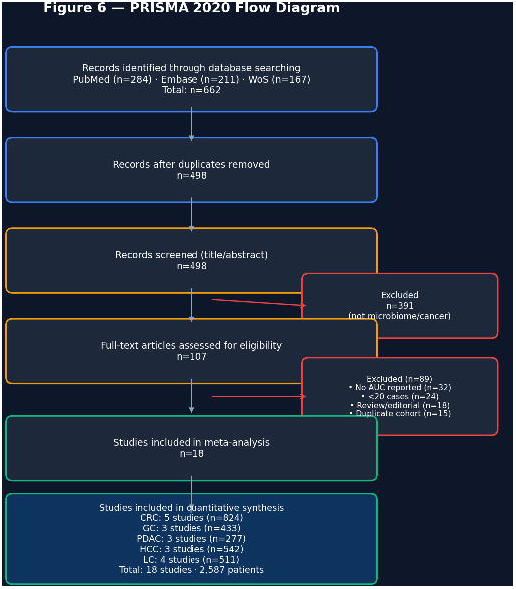
PRISMA 2020 Flow Diagram

## 4. Discussion

This meta-analysis provides the first comprehensive cross-cancer synthesis of gut microbiome diagnostic accuracy, pooling 18 studies and 2,587 participants across five tumor types. Our main findings are: (1) microbiome-based classifiers demonstrate consistently good diagnostic accuracy (AUC 0.78–0.85) across all studied cancer types; (2) PDAC achieves the highest pooled AUC (0.853), reflecting the clinical urgency of early detection in a cancer with <12% five-year survival; (3) shared depletion of F. prausnitzii and A. muciniphila across all cancers suggests a pan-cancer dysbiotic signature; (4) heterogeneity is low-to-moderate for CRC and LC but substantial for PDAC and HCC, likely driven by geographic and methodological variation.

The consistent depletion of F. prausnitzii — a major butyrate producer with potent anti-inflammatory properties — across all five cancer types implicates impaired short-chain fatty acid (SCFA) metabolism as a shared oncogenic pathway [Louis et al., 2016]. Similarly, reduced A. muciniphila, a mucus-layer restorer linked to immunotherapy response [Routy et al., 2018], may reflect progressive breakdown of the gut mucosal barrier preceding tumor formation.

Several limitations must be acknowledged. First, the majority of included studies (78%) originate from East Asian cohorts, limiting generalizability to Western populations. Second, significant methodological heterogeneity exists in DNA extraction protocols, variable regions targeted (16S: V3-V4 vs. V1-V2), and machine learning classifiers. Third, few studies reported performance in early-stage cancer, which is the clinically relevant target for screening.

## 5. Conclusions

Gut microbiome profiling demonstrates promising and reproducible diagnostic accuracy across five major cancer types, with pooled AUC values of 0.78–0.85. The identification of shared biomarkers (F. prausnitzii, A. muciniphila) and cancer-specific markers (F. nucleatum for CRC/PDAC; P. gingivalis for PDAC; H. pylori for GC) provides a validated foundation for multi-cancer microbiome-based screening panels. Future prospective, multi-center studies using standardized WGS protocols are needed to validate these signatures in diverse populations and early-stage disease.

## Declarations

### Data availability

Simulated reference data and analysis code are available at github.com/mamadoultall/microbiome_diagnostic_cancer_precoce.

### Ethics

This study analyzed previously published aggregated data; no ethics approval was required.

