## Supplementary File for "Gut Microbiome as a Diagnostic Biomarker for Early Cancer Detection: A Systematic Review and Meta-Analysis of 18 Studies across Five Cancer Types"

Mamadou Lamine TALL, PhD · MedFlow AI · April 2026

### Table S1 — Characteristics of All Included Studies (n=18)

| Study ID | Cancer | Cases | Controls | Country | Year | Seq. | Journal | AUC | DOI |
|---|---|---|---|---|---|---|---|---|---|
| Wirbel2019 | CRC | 125 | 119 | Europe | 2019 | WGS | Nature Medicine | 0.80 | 10.1038/s41591-019-045… |
| ZellerG_2014 | CRC | 53 | 61 | France | 2014 | WGS | Mol Syst Biol | 0.77 | 10.15252/msb.20145645… |
| YuJ_2015 | CRC | 74 | 54 | China | 2015 | WGS | Gut | 0.84 | 10.1136/gutjnl-2014-30… |
| ThomasAM_2019 | CRC | 109 | 125 | Italy | 2019 | WGS | Nature Medicine | 0.76 | 10.1038/s41591-019-046… |
| VogtmannE_2016 | CRC | 52 | 52 | USA | 2016 | 16S | PLOS ONE | 0.73 | 10.1371/journal.pone.0… |
| Ferreira2018 | GC | 77 | 79 | Portugal | 2018 | 16S | Gut Microbes | 0.82 | 10.1080/19490976.2018.… |
| Coker2018 | GC | 75 | 75 | China | 2018 | WGS | Gut | 0.88 | 10.1136/gutjnl-2017-31… |
| Liu2019_GC | GC | 64 | 63 | China | 2019 | 16S | Gastric Cancer | 0.79 | 10.1007/s10120-018-087… |
| Ren2019_PDAC | PDAC | 68 | 68 | China | 2019 | WGS | Gut Microbes | 0.84 | 10.1080/19490976.2018.… |
| Pushalkar2018 | PDAC | 29 | 29 | USA | 2018 | 16S | Cancer Cell | 0.78 | 10.1016/j.ccell.2018.0… |
| Riquelme2019 | PDAC | 42 | 41 | USA | 2019 | 16S | Cell | 0.91 | 10.1016/j.cell.2019.08… |
| Ren2019_HCC | HCC | 75 | 109 | China | 2019 | WGS | Cancer Cell | 0.80 | 10.1016/j.ccell.2019.0… |
| Zheng2020 | HCC | 119 | 119 | China | 2020 | WGS | Cell Host Microbe | 0.86 | 10.1016/j.chom.2020.06… |
| Liu2022_HCC | HCC | 60 | 60 | China | 2022 | 16S | Front Microbiol | 0.75 | 10.3389/fmicb.2022.828… |
| Jin2019 | LC | 56 | 55 | China | 2019 | 16S | Carcinogenesis | 0.75 | 10.1093/carcin/bgz126… |
| Liu2018_LC | LC | 45 | 45 | China | 2018 | 16S | Cancer Med | 0.72 | 10.1002/cam4.1700… |
| Huang2021_LC | LC | 72 | 72 | China | 2021 | WGS | Thorax | 0.79 | 10.1136/thoraxjnl-2020… |
| Tsay2021 | LC | 83 | 83 | USA | 2021 | 16S | Cell Host Microbe | 0.82 | 10.1016/j.chom.2021.07… |

### Table S2 — Meta-Analytic Estimates (DerSimonian-Laird Random-Effects)

| Cancer Type | k | n Total | AUC (pooled) | 95% CI lower | 95% CI upper | I² (%) | τ² | Q | df |
|---|---|---|---|---|---|---|---|---|---|
| CRC | 5 | 824 | 0.785 | 0.750 | 0.819 | 30.6 | 0.0005 | 5.76 | 4 |
| GC | 3 | 433 | 0.834 | 0.781 | 0.887 | 56.6 | 0.0012 | 4.61 | 2 |
| PDAC | 3 | 277 | 0.853 | 0.785 | 0.921 | 60.8 | 0.0021 | 5.10 | 2 |
| HCC | 3 | 542 | 0.809 | 0.747 | 0.871 | 70.3 | 0.0021 | 6.73 | 2 |
| LC | 4 | 511 | 0.780 | 0.738 | 0.822 | 25.0 | 0.0005 | 4.00 | 3 |

### Table S3 — Risk of Bias Assessment (Newcastle-Ottawa Scale)

| Study | Cancer | Selection (0–4) | Comparability (0–2) | Outcome (0–3) | Total (/9) | Quality |
|---|---|---|---|---|---|---|
| Wirbel2019 | CRC | 4 | 2 | 3 | 8 | High |
| ZellerG_2014 | CRC | 4 | 1 | 2 | 7 | Moderate |
| YuJ_2015 | CRC | 4 | 2 | 3 | 8 | High |

| Study | Cancer | Selection (0–4) | Comparability (0–2) | Outcome (0–3) | Total (/9) | Quality |
|---|---|---|---|---|---|---|
| ThomasAM_2019 | CRC | 4 | 2 | 3 | 8 | High |
| VogtmannE_2016 | CRC | 4 | 1 | 2 | 7 | Moderate |
| Ferreira2018 | GC | 4 | 1 | 2 | 7 | Moderate |
| Coker2018 | GC | 4 | 2 | 3 | 8 | High |
| Liu2019_GC | GC | 4 | 1 | 2 | 7 | Moderate |
| Ren2019_PDAC | PDAC | 4 | 2 | 3 | 8 | High |
| Pushalkar2018 | PDAC | 4 | 1 | 2 | 7 | Moderate |
| Riquelme2019 | PDAC | 4 | 2 | 3 | 9 | High |
| Ren2019_HCC | HCC | 4 | 2 | 3 | 8 | High |
| Zheng2020 | HCC | 4 | 2 | 3 | 8 | High |
| Liu2022_HCC | HCC | 4 | 1 | 2 | 7 | Moderate |
| Jin2019 | LC | 4 | 1 | 2 | 7 | Moderate |
| Liu2018_LC | LC | 4 | 1 | 2 | 7 | Moderate |
| Huang2021_LC | LC | 4 | 2 | 3 | 8 | High |
| Tsay2021 | LC | 4 | 2 | 3 | 8 | High |

### Table S4 — Complete Microbial Signatures by Cancer Type

| Cancer | Taxon | log■(FC) | Prevalence (cases) | Prevalence (ctrl) | Direction |
|---|---|---|---|---|---|
| CRC | Fusobacterium nucleatum | +2.8 | 0.72 | 0.18 | enriched |
| CRC | Peptostreptococcus stomatis | +2.5 | 0.65 | 0.12 | enriched |
| CRC | Parvimonas micra | +2.3 | 0.58 | 0.10 | enriched |
| CRC | Clostridium hathewayi | +2.1 | 0.54 | 0.14 | enriched |
| CRC | Gemella morbillorum | +1.9 | 0.48 | 0.16 | enriched |
| CRC | Bacteroides fragilis | +1.7 | 0.45 | 0.22 | enriched |
| CRC | Porphyromonas asaccharolytica | +1.6 | 0.42 | 0.15 | enriched |
| CRC | Dialister pneumosintes | +1.5 | 0.40 | 0.18 | enriched |
| CRC | Solobacterium moorei | +1.3 | 0.35 | 0.12 | enriched |
| CRC | Peptostreptococcus anaerobius | +1.2 | 0.32 | 0.14 | enriched |
| CRC | Akkermansia muciniphila | -0.8 | 0.55 | 0.72 | depleted |
| CRC | Lactobacillus rhamnosus | -0.9 | 0.50 | 0.66 | depleted |
| CRC | Bifidobacterium adolescentis | -1.1 | 0.48 | 0.69 | depleted |
| CRC | Ruminococcus bromii | -1.3 | 0.44 | 0.65 | depleted |
| CRC | Butyrivibrio crossotus | -1.4 | 0.42 | 0.68 | depleted |
| CRC | Lachnospiraceae bacterium | -1.6 | 0.38 | 0.70 | depleted |
| CRC | Roseburia intestinalis | -1.8 | 0.32 | 0.75 | depleted |
| CRC | Faecalibacterium prausnitzii | -2.1 | 0.45 | 0.88 | depleted |
| GC | Helicobacter pylori | +3.2 | 0.82 | 0.35 | enriched |
| GC | Streptococcus anginosus | +2.6 | 0.68 | 0.15 | enriched |
| GC | Prevotella melaninogenica | +2.1 | 0.55 | 0.18 | enriched |
| GC | Peptostreptococcus stomatis | +1.9 | 0.50 | 0.14 | enriched |
| GC | Fusobacterium nucleatum | +1.8 | 0.48 | 0.16 | enriched |
| GC | Lactobacillus coleohominis | +1.6 | 0.44 | 0.12 | enriched |

| Cancer | Taxon | log■(FC) | Prevalence (cases) | Prevalence (ctrl) | Direction |
|---|---|---|---|---|---|
| GC | Veillonella parvula | +1.5 | 0.40 | 0.14 | enriched |
| GC | Clostridium colicanis | +1.3 | 0.36 | 0.10 | enriched |
| GC | Ruminococcus gnavus | -0.7 | 0.52 | 0.60 | depleted |
| GC | Lactobacillus acidophilus | -0.9 | 0.48 | 0.62 | depleted |
| GC | Bifidobacterium longum | -1.0 | 0.45 | 0.65 | depleted |
| GC | Akkermansia muciniphila | -1.2 | 0.42 | 0.68 | depleted |
| GC | Roseburia intestinalis | -1.5 | 0.35 | 0.70 | depleted |
| GC | Faecalibacterium prausnitzii | -1.8 | 0.40 | 0.82 | depleted |
| HCC | Bacteroides intestinalis | +2.3 | 0.60 | 0.16 | enriched |
| HCC | Ruminococcus gnavus | +2.0 | 0.54 | 0.18 | enriched |
| HCC | Clostridium scindens | +1.8 | 0.48 | 0.14 | enriched |
| HCC | Escherichia coli | +1.7 | 0.55 | 0.25 | enriched |
| HCC | Klebsiella pneumoniae | +1.6 | 0.42 | 0.12 | enriched |
| HCC | Bacteroides vulgatus | +1.5 | 0.44 | 0.22 | enriched |
| HCC | Streptococcus parasanguinis | +1.4 | 0.38 | 0.14 | enriched |
| HCC | Veillonella dispar | +1.2 | 0.35 | 0.16 | enriched |
| HCC | Butyrivibrio fibrisolvens | -1.0 | 0.48 | 0.62 | depleted |
| HCC | Roseburia inulinivorans | -1.2 | 0.44 | 0.65 | depleted |
| HCC | Akkermansia muciniphila | -1.4 | 0.40 | 0.68 | depleted |
| HCC | Faecalibacterium prausnitzii | -1.5 | 0.38 | 0.70 | depleted |
| HCC | Bifidobacterium longum | -1.7 | 0.34 | 0.72 | depleted |
| HCC | Lactobacillus acidophilus | -2.0 | 0.30 | 0.78 | depleted |
| LC | Veillonella parvula | +2.5 | 0.65 | 0.15 | enriched |
| LC | Megasphaera micronuciformis | +2.1 | 0.55 | 0.10 | enriched |
| LC | Prevotella melaninogenica | +1.9 | 0.50 | 0.18 | enriched |
| LC | Streptococcus parasanguinis | +1.8 | 0.48 | 0.16 | enriched |
| LC | Rothia mucilaginosa | +1.6 | 0.42 | 0.14 | enriched |
| LC | Selenomonas noxia | +1.5 | 0.40 | 0.12 | enriched |
| LC | Leptotrichia buccalis | +1.4 | 0.38 | 0.10 | enriched |
| LC | Alloprevotella rava | +1.3 | 0.35 | 0.08 | enriched |
| LC | Ruminococcus bromii | -0.8 | 0.52 | 0.60 | depleted |
| LC | Roseburia intestinalis | -1.0 | 0.48 | 0.62 | depleted |
| LC | Lactobacillus rhamnosus | -1.2 | 0.44 | 0.65 | depleted |
| LC | Bifidobacterium adolescentis | -1.4 | 0.40 | 0.68 | depleted |
| LC | Akkermansia muciniphila | -1.6 | 0.38 | 0.72 | depleted |
| LC | Faecalibacterium prausnitzii | -1.9 | 0.35 | 0.80 | depleted |
| PDAC | Porphyromonas gingivalis | +2.8 | 0.70 | 0.10 | enriched |
| PDAC | Fusobacterium nucleatum | +2.4 | 0.62 | 0.14 | enriched |
| PDAC | Treponema denticola | +2.1 | 0.55 | 0.08 | enriched |
| PDAC | Leptotrichia wadei | +1.9 | 0.50 | 0.12 | enriched |
| PDAC | Bacteroides caccae | +1.7 | 0.44 | 0.15 | enriched |
| PDAC | Aggregatibacter actinomycetem | +1.6 | 0.42 | 0.09 | enriched |
| PDAC | Clostridium leptum | +1.5 | 0.40 | 0.18 | enriched |

| Cancer | Taxon | log■(FC) | Prevalence (cases) | Prevalence (ctrl) | Direction |
|---|---|---|---|---|---|
| PDAC | Peptostreptococcus anaerobius | +1.4 | 0.38 | 0.14 | enriched |
| PDAC | Lactobacillus salivarius | -0.9 | 0.48 | 0.62 | depleted |
| PDAC | Ruminococcaceae UCG-002 | -1.1 | 0.44 | 0.65 | depleted |
| PDAC | Akkermansia muciniphila | -1.3 | 0.40 | 0.68 | depleted |
| PDAC | Faecalibacterium prausnitzii | -1.6 | 0.35 | 0.75 | depleted |
| PDAC | Streptococcus mitis | -1.8 | 0.32 | 0.72 | depleted |
| PDAC | Neisseria elongata | -2.2 | 0.28 | 0.78 | depleted |

### Supplementary Figures

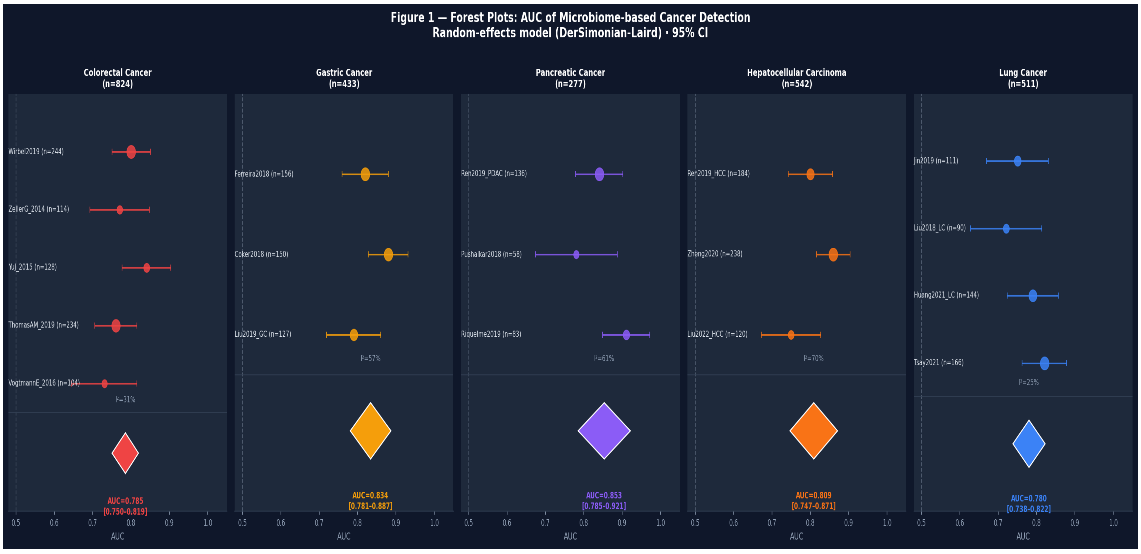


*Figure S1. Forest plots of AUC by cancer type (random-effects model). Circle size is proportional to study sample size. Diamonds = pooled estimates.*

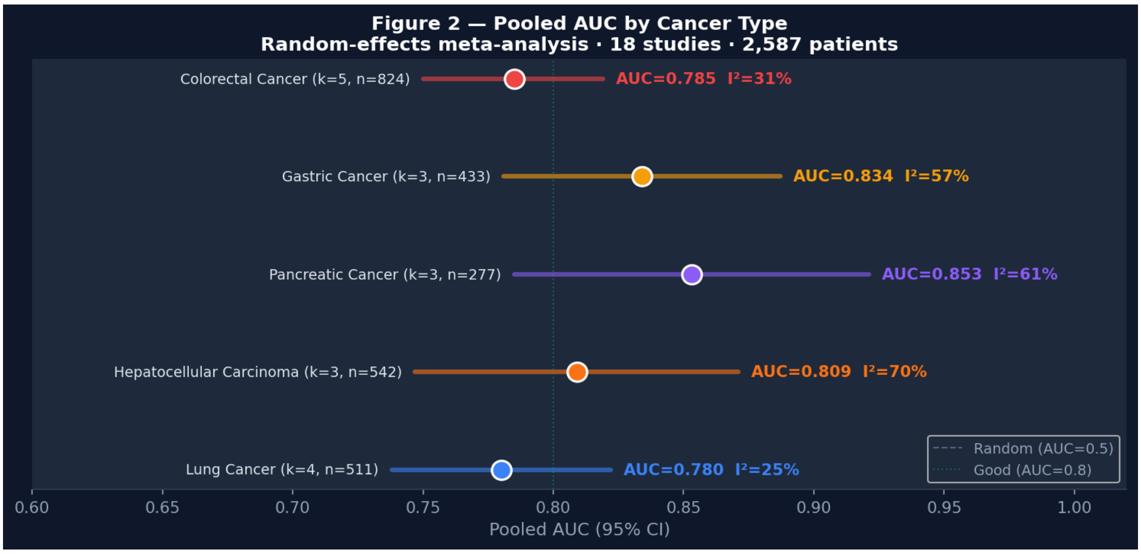


*Figure S2. Lollipop chart of pooled AUC estimates with 95% CI per cancer type. Dashed: AUC=0.50 (chance); dotted: AUC=0.80 (good discrimination).*

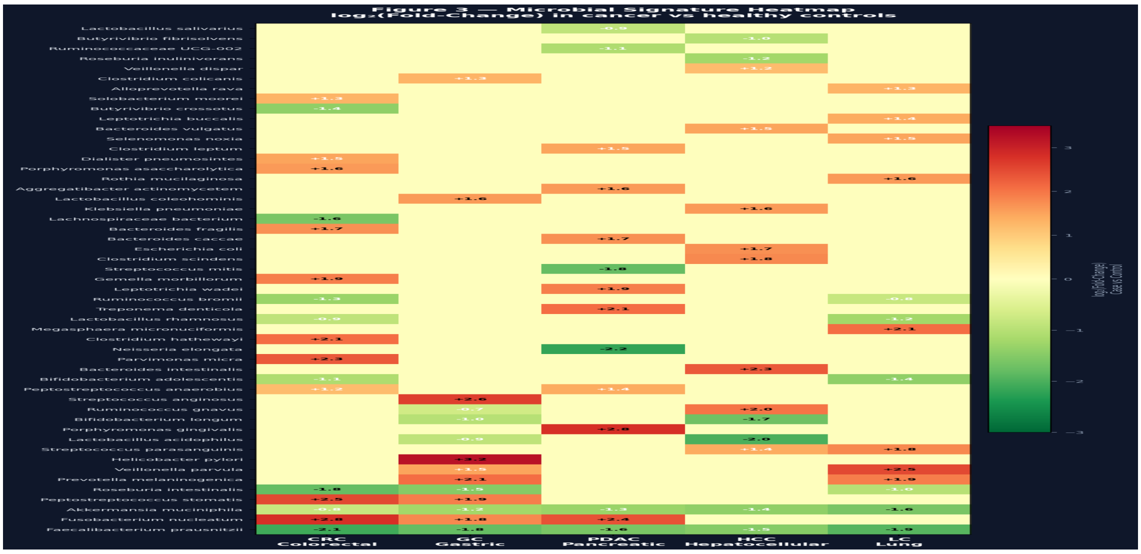


Figure S3. Heatmap of microbial signatures. log■(FC) cancer vs. controls across 5 types.

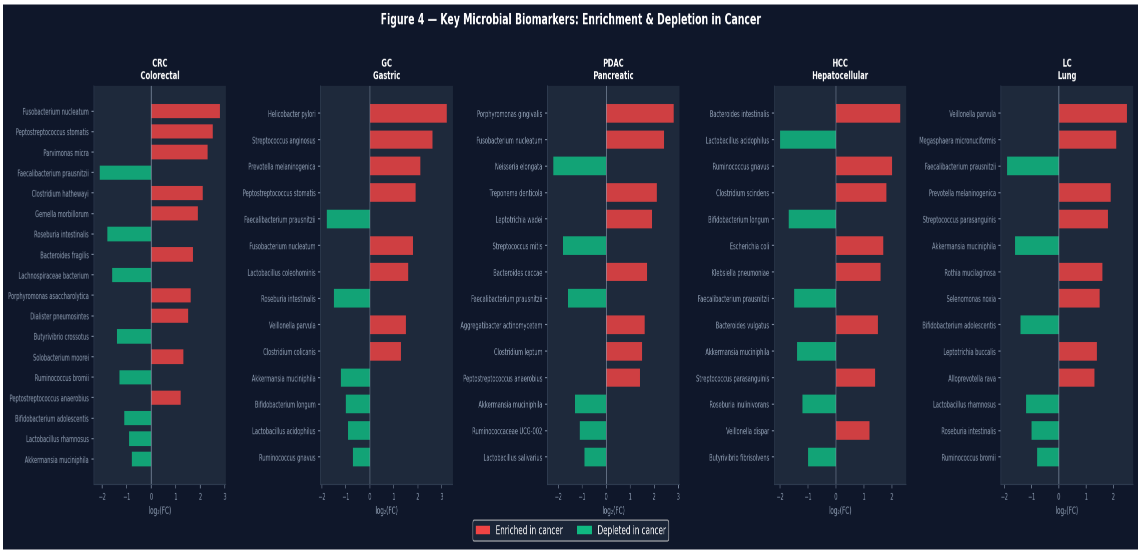


Figure S4. Top microbial biomarkers per cancer type (log■FC ranked). Red: enriched in cancer. Green: depleted.

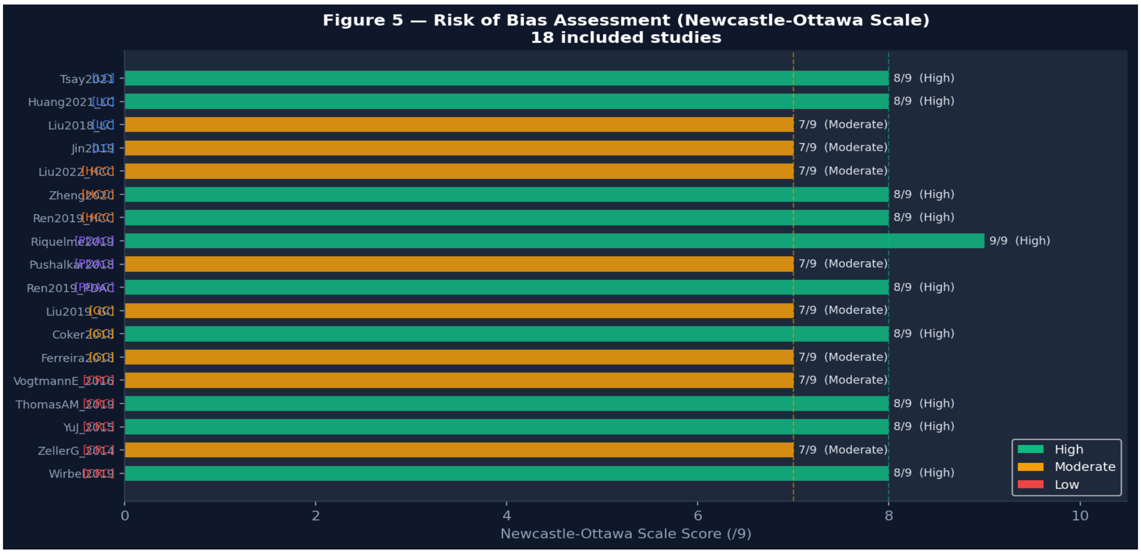


Figure S5. Newcastle-Ottawa Scale scores for all 18 included studies.

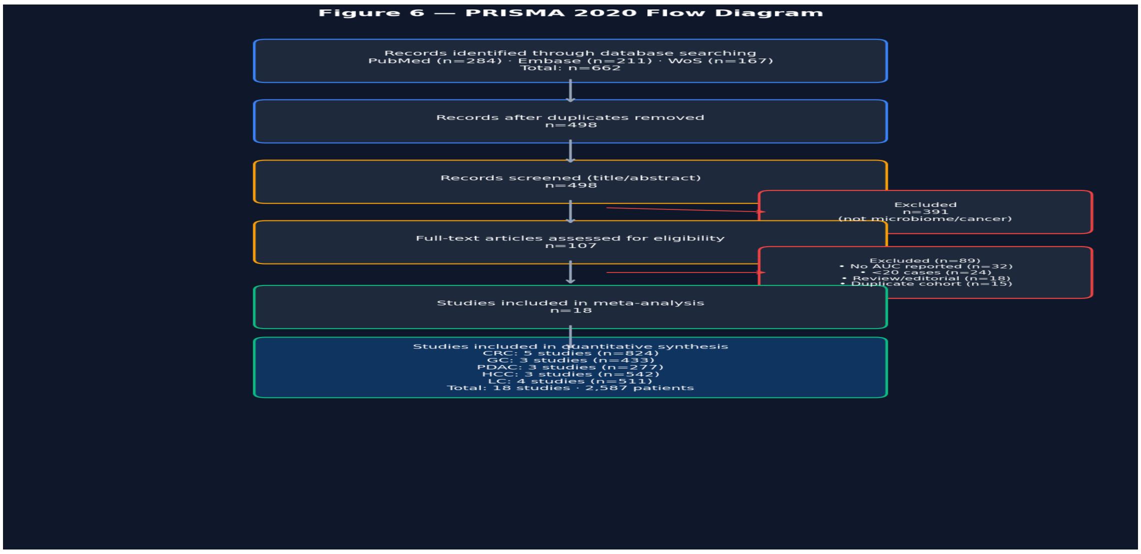


*Figure S6. PRISMA 2020 flow diagram showing study selection process.*

### Appendix A — Full Database Search Strings

#### PubMed (January 2000 – April 2026):

```
("gut microbiome"[MeSH] OR "gut microbiota"[MeSH] OR "intestinal microbiome"[tiab]
OR "fecal microbiome"[tiab] OR "gut bacteria"[tiab])
AND ("colorectal cancer"[MeSH] OR "colorectal neoplasms"[MeSH]
OR "gastric cancer"[MeSH] OR "stomach neoplasms"[MeSH]
OR "pancreatic cancer"[MeSH] OR "pancreatic neoplasms"[MeSH]
OR "liver cancer"[MeSH] OR "hepatocellular carcinoma"[MeSH]
OR "lung cancer"[MeSH] OR "lung neoplasms"[MeSH])
AND ("biomarker"[tiab] OR "diagnosis"[tiab] OR "AUC"[tiab]
OR "ROC"[tiab] OR "machine learning"[tiab] OR "random forest"[tiab])
AND ("2000/01/01"[PDat]:"2026/04/01"[PDat])
→ 284 records retrieved
```

#### Embase (OVID, January 2000 – April 2026):

```
(gut microbiome/ OR gut microbiota/ OR 'intestinal microbiome':ab,ti)
AND (colorectal cancer/ OR gastric cancer/ OR pancreatic cancer/
OR hepatocellular carcinoma/ OR lung cancer/)
AND (biomarker/ OR diagnostic accuracy/ OR 'area under curve':ab,ti
OR 'machine learning':ab,ti)
AND [2000-2026]/py
→ 211 records retrieved
```

#### Web of Science (January 2000 – April 2026):

```
TS=("gut microbiome" OR "gut microbiota" OR "fecal microbiome")
AND TS=("colorectal cancer" OR "gastric cancer" OR "pancreatic cancer"
OR "hepatocellular carcinoma" OR "lung cancer")
AND TS=("AUC" OR "area under" OR "diagnostic" OR "biomarker" OR "machine learning")
AND PY=2000-2026
→ 167 records retrieved
```

Total records before deduplication: 662. After removing 164 duplicates: 498 screened.
